# Associations between Afrotropical bats, parasites, and microbial symbionts

**DOI:** 10.1101/340109

**Authors:** L. Lutz Holly, W. Jackson Elliot, W. Dick Carl, W. Webala Paul, S. Babyesiza Waswa, C. Kerbis Peterhans Julian, C. Demos Terrence, D. Patterson Bruce, A. Gilbert Jack

**Author notes:** Corresponding author: Holly L. Lutz & Jack A.Gilbert, Department of Surgery, Division of the Biological Sciences, University of Chicago, 5851 S. Maryland Avenue Chicago, IL 60637 - 1508.

## Abstract

Bats are among the most diverse animals on the planet and harbor numerous bacterial, viral, and eukaryotic symbionts. The interplay between bacterial community composition and parasitism in bats is not well understood and may have important implications for studies of similar systems. Here we present a comprehensive survey of dipteran and haemosporidian parasites, and characterize the gut, oral, and skin microbiota of Afrotropical bats. We identify significant correlations between bacterial community composition of the skin and dipteran ectoparasite prevalence across four major bat lineages, as well as links between the oral microbiome and malarial parasitism, suggesting a potential mechanism for host selection and vector-borne disease transmission in bats. In contrast to recent studies of host-microbe phylosymbiosis in mammals, we find no correlation between chiropteran phylogenetic distances and bacterial community dissimilarity across the three anatomical sites, suggesting that host environment is more important than shared ancestry in shaping the composition of bat-associated bacterial communities.

**SIGNIFICANCE:** Animals rely on bacterial symbionts for numerous biological functions, such as digestion and immune system development. Increasing evidence suggests that host-associated microbes may play a role in mediating parasite burden. This study is the first to provide a comprehensive survey of bacterial symbionts from multiple anatomical sites across a broad taxonomic range of Afrotropical bats, demonstrating significant associations between the bat microbiome and parasite prevalence. This study provides a framework for future approaches to systems biology of host-symbiont interactions across broad taxonomic scales, emphasizing the interdependence between microbial symbionts and vertebrate health in the study of wild organisms and their natural history.

## INTRODUCTION

Humans and other animals rely on bacterial symbionts for numerous biological functions, such as digestion and immune system development (1, 2). Many studies have found significant associations between host phylogeny (shared common ancestry) and bacterial community composition (3, 4), while others have identified dietary or spatiotemporal variables as significant drivers of host-microbe associations over the course of an individual lifespan (5-7). The influence of microbes on their hosts may be context dependent, such that the presence of a particular microbe may be beneficial under one set of ecological conditions and harmful under another. Thus, patterns of association between animals and bacterial symbionts provide a unique lens through which to explore evolutionary and ecological phenomena.

Recognition of the interdependence between microbial symbionts and animal health has led to a growing paradigm shift in the study of wild organisms and their natural history. In addition to exhibiting variation in life history characteristics, animals serve as hosts to myriad bacteria, archaea, viruses, fungi, and eukaryotic organisms. Many relationships between eukaryotic parasites and hosts have ancient origins, and the same may be true for host-microbial associations. It is possible that bacterial symbionts of vertebrate hosts interact with eukaryotic parasites, viruses, or fungal symbionts in ways that could shape host evolution (8). For example, evidence from human and anthropophilic mosquito interactions suggests that the skin microbiome can influence vector feeding preference, thereby affecting transmission patterns of mosquito-borne pathogens (such as West Nile virus, yellow fever, dengue, malaria, etc.), and ultimately imposing selective pressures on human populations - indeed, positive selection of malaria-protective genes can be seen in the human genome (9). Despite the potential significance of such interactions between hosts, microbes, and pathogen-transmitting vectors, they have not been well studied in most wild vertebrate systems.

Bats (Mammalia: Chiroptera) are an important system for comparison of the relative contributions of evolutionary and ecological factors driving host-symbiont associations. In addition to being one of the most speciose orders of mammals (second only to the order Rodentia), bats frequently live in large colonies, are long-lived, and volant, granting them access to a wide geographic range relative to their non-volant mammalian counterparts. The associations of diverse eukaryotic parasites (*e.g.* dipteran insects, haemosporidia, helminths) within numerous bat lineages have been well-characterized (10-13). Furthermore, bats have received increasing attention due to their role as reservoirs of human pathogens (*e.g.* Ebola, Marburg, Nipah, SARS (14-18)). Taken together, these features make bats an important and tractable model for studying the interaction of bacterial symbionts and non-bacterial parasites and pathogens.

In this study, we conduct the first broad-scale study of Afrotropical bat-associated microbes. We test associations between bacterial community composition in the gastrointestinal tract, skin, and oral cavities from nine families and nineteen genera of bats. We pair this information with host-parasite associations between bats and ectoparasites in the superfamily Hippoboscoidea (obligate hematophagous dipteran insects), and haemosporidian (malarial) parasites putatively vectored by these hippoboscoid insects. Using a combination of machine learning, network theory, and negative binomial distribution models, we test the hypothesis that host-associated bacterial communities predict prevalence of parasitism by obligate dipteran and malarial parasites.

### RESULTS

### 1) Ectoparasite and malarial parasite prevalence among Afrotropical bats

Sampling was conducted across 20 sites in Kenya and Uganda from July-August of 2016. Sites ranged from sea level to ∼2500m in elevation (Fig. 1; Table S1). We collected gut, oral, and skin samples for bacterial community characterization from a total of 494 individual bats, comprising 9 families, 19 genera, and 28 recognized species. Bat families with the greatest representation included Hipposideridae (*n* = 80), Miniopteridae (*n* = 116), Rhinolophidae (*n* = 88), and Pteropodidae (*n* = 106). All host and parasite vouchers are accessioned at the Field Museum of Natural History (Chicago, IL, USA) (Table S2). Miniopterid bats experienced the highest prevalence of both ectoparasitism (*M. minor*, 89%) and malarial parasitism (*M. minor*, 67%) (Table 1). Bats with similarly high ectoparasite prevalence at the host species level included *Rhinolophus eloquens* (79% prevalence), *Stenonycteris lanosus* (62%), and *Triaenops afer* (60%). Unlike miniopterid bats, these bat species did not harbor any detectable malarial parasites (Table 1).

**Table 1.**
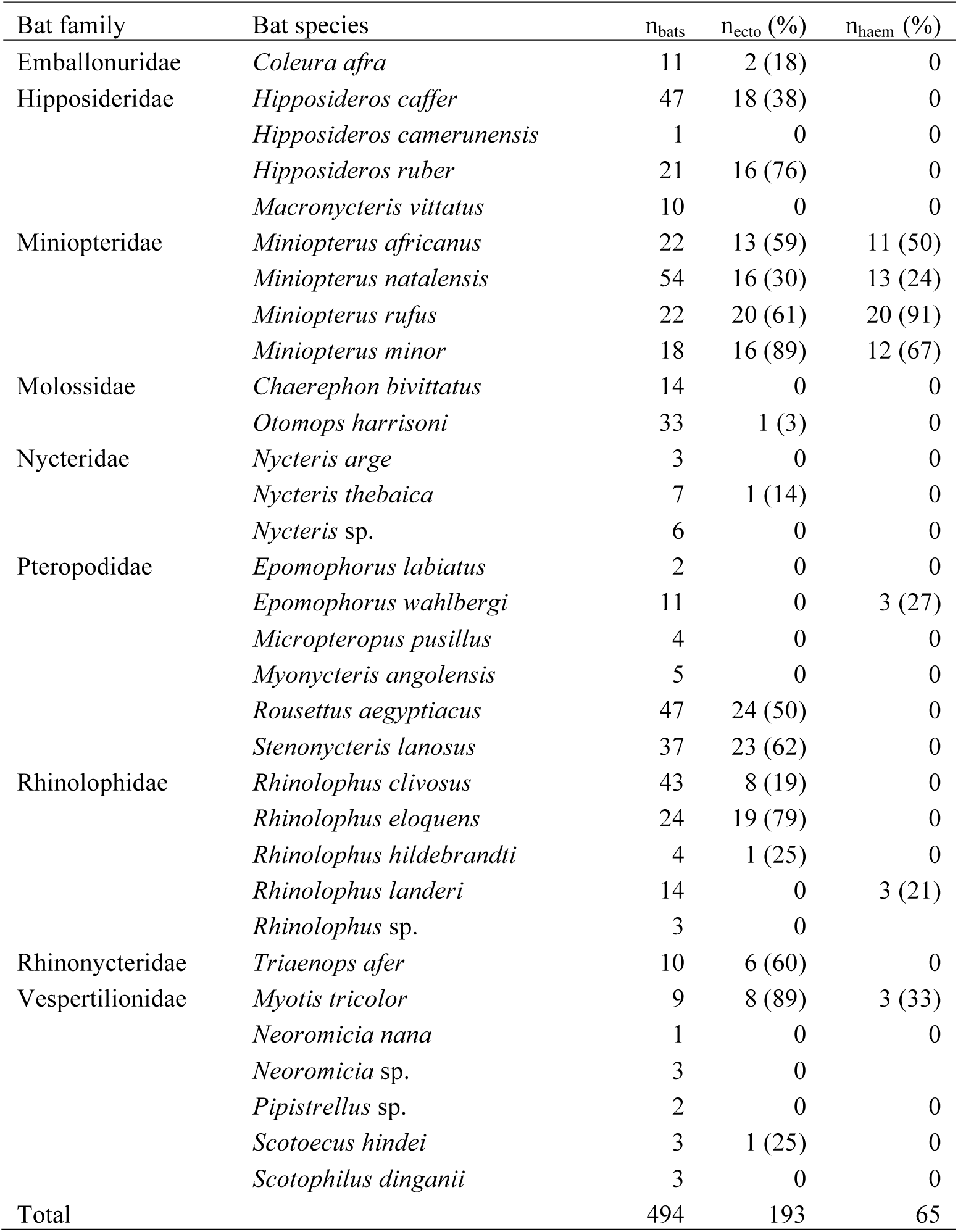
Bat sampling, ectoparasite prevalence (n_ecto_), and malarial parasite prevalence (n_haem_) and identification.

**Figure 1.**
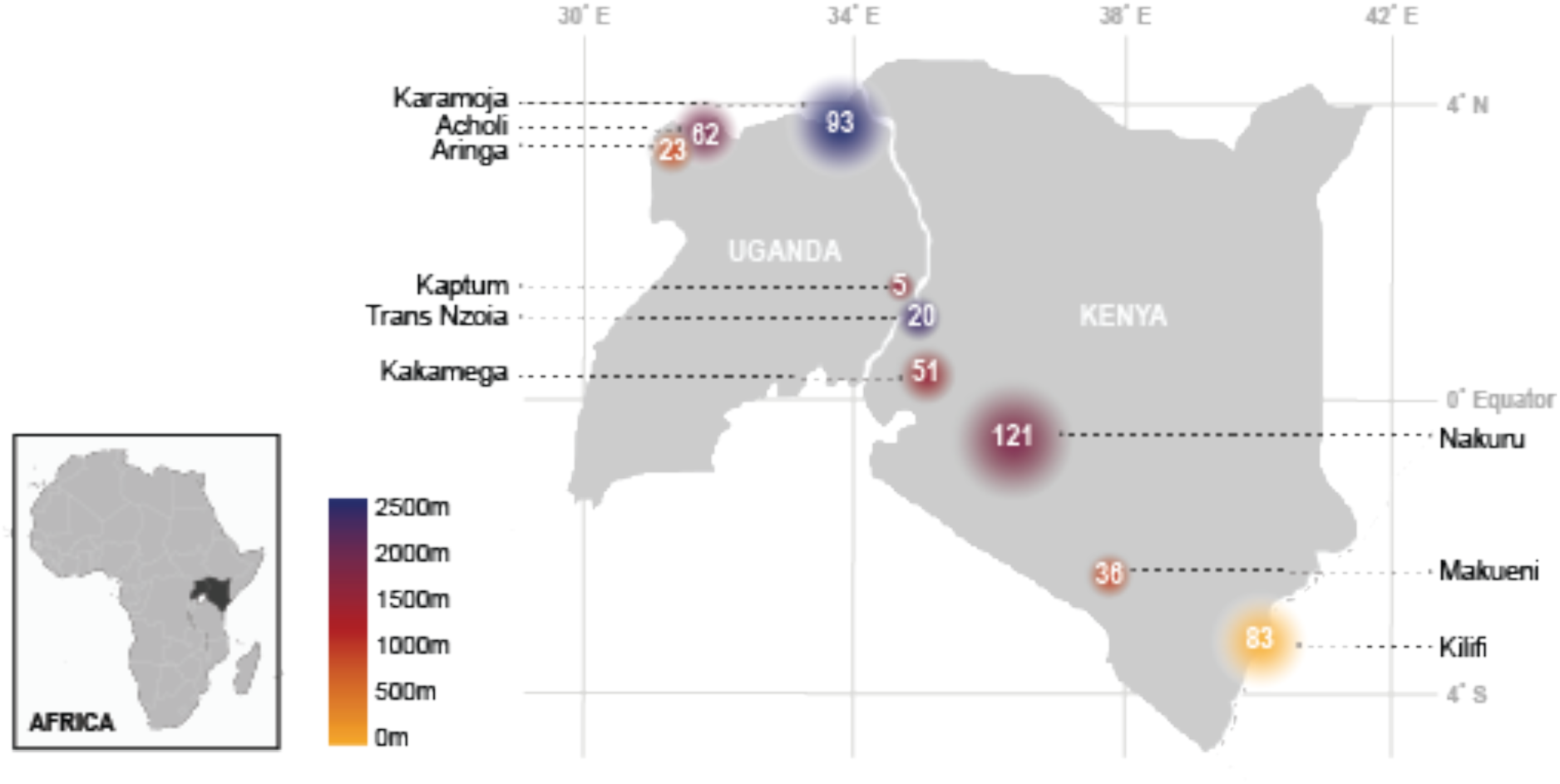
Sampling localities and elevation, grouped by county (see Table S1 for full locality information). Colors correspond to elevation, and white numbers and size of points correspond to number of bats collected.

### 2 Microbial richness associated with bat skin is significantly greater than gut or oral microbial communities

Sequencing produced a total of 1,236 libraries, with an average read depth of 32,950 reads per library (±19,850 reads). Analyses and statistical tests were performed on non-rarefied data (libraries containing >1000 reads and transformed to library read depth) and on rarefied data (libraries rarefied to a read depth of 10,000 sequences and subsequently transformed). The difference in library normalization methods only resulted in a decrease of total ASVs which did not affect the significance of alpha and beta-diversity statistical tests. Therefore, the results reported hereafter are from the non-rarefied data set. Across all samples, 31,172 amplicon sequence variants (ASVs) were identified using Deblur (19) (for rarefied data set, 1,267 ASVs were dropped, resulting in a total of 29,890 ASVs identified across all samples). Total number of libraries per anatomical site, following filtering, included 396 libraries for gut, 374 libraries for oral, and 458 libraries for skin microbiomes (Table 2). Gut microbial communities exhibited the lowest overall diversity (8,204 ASVs), followed by oral (11,632 ASVs), and skin (29,149 ASVs), the latter being significantly greater than gut or oral (*p* < 2.2e-16, Kruskal-Wallis; Bonferroni corrected *p*-value *p* < 1e-113, Dunn’s test) (Fig. 2; Fig S1). The mean observed ASVs by anatomical site were 69, 96, and 587 for gut, oral, and skin samples, respectively (Table 2). Shannon index score of skin microbial communities were also significantly greater than gut or oral microbial communities (*p* < 2.2e-16, Kruskal-Wallis; Bonferroni corrected *p*-value *p* < 1e-119, Dunn’s Test) (Fig. 2; Fig. S1).

**Table 2.**
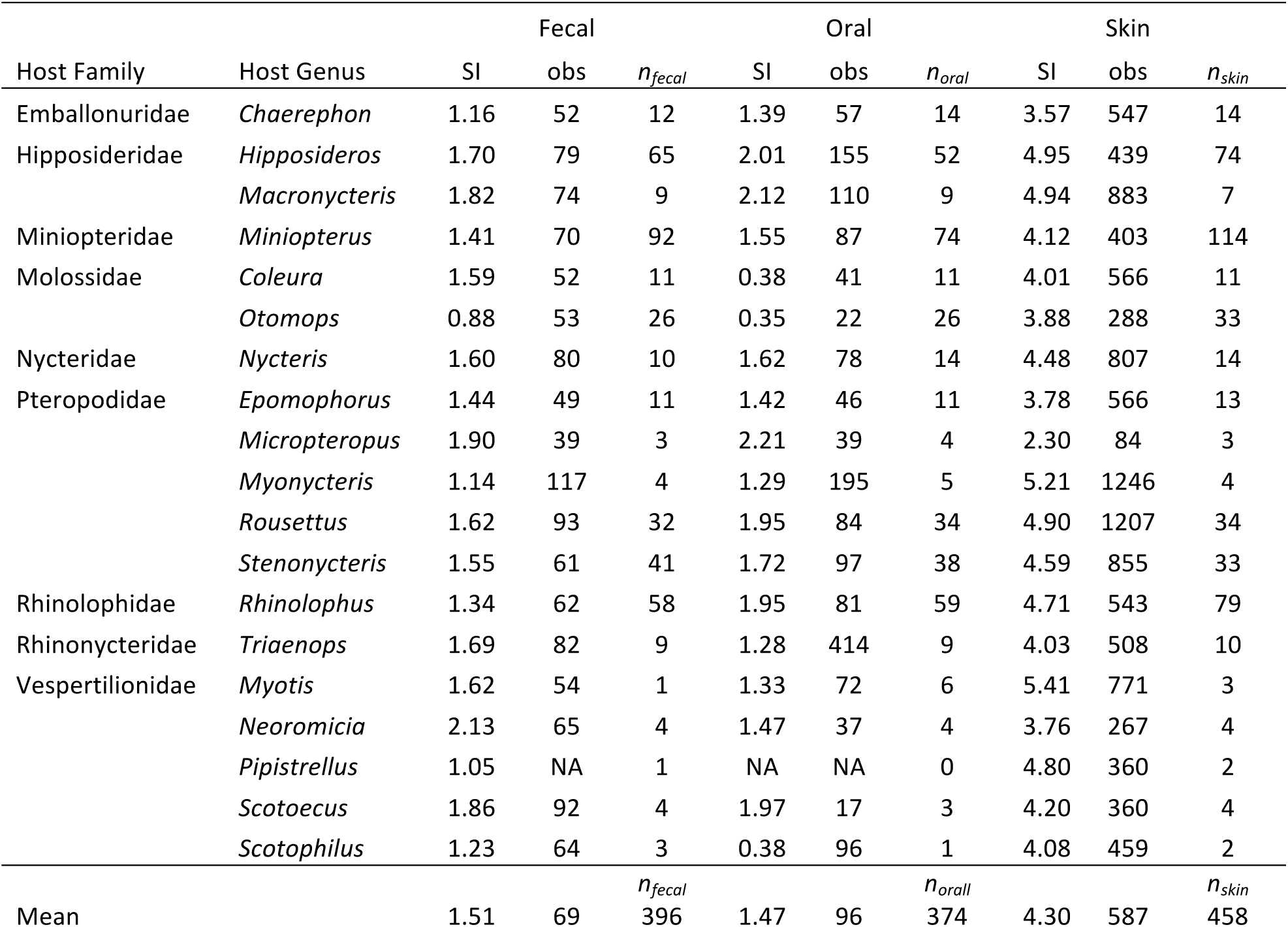
Alpha diversity of microbial communities across anatomical sites within each host genus, measured by Shannon Index of diversity (SI) and observed ASV richness (obs); n corresponds to number of libraries included in each calculation (following quality filtering).

**Figure 2.**
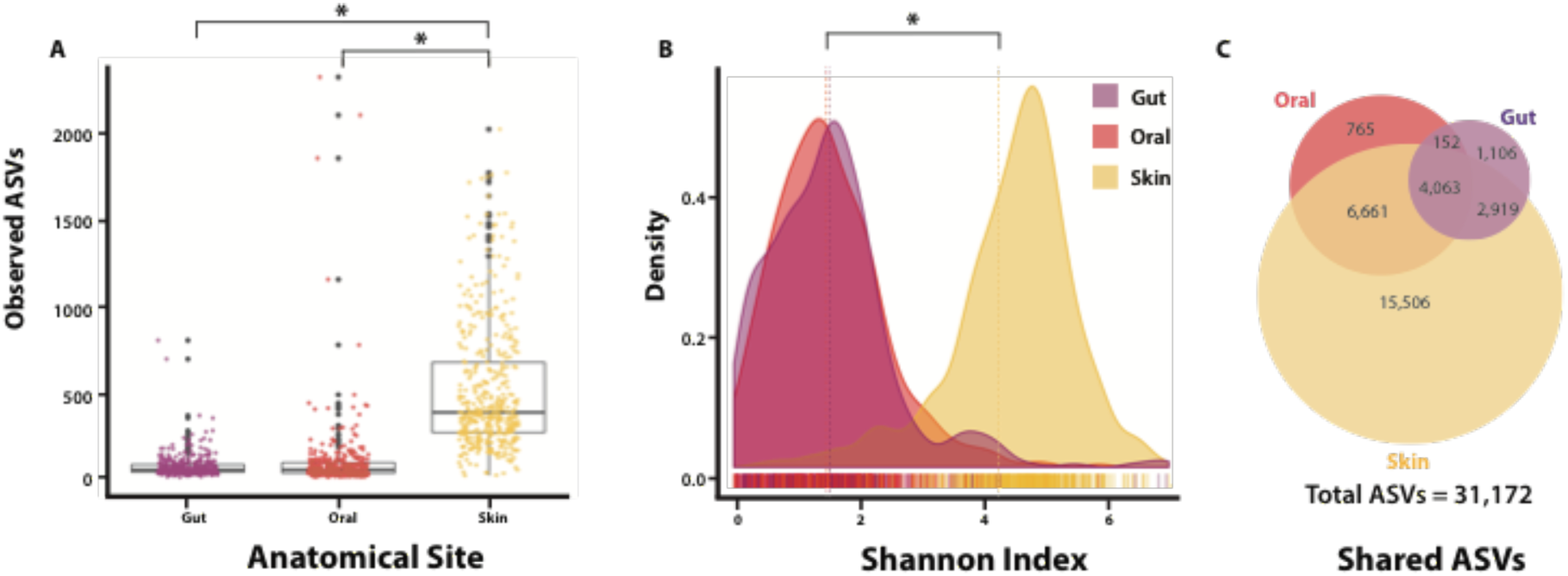
Alpha diversity of amplicon sequence variants (ASVs) by anatomical sites, including (A) Observed richness, (B) Shannon index of diversity, (C) ASVs shared between anatomical sites. Asterisks indicate significant differences between groups (Dunn’s Test, Bonferroni corrected *p*-value *p* < 0.0001).

### 3 Microbial communities significantly correlate with geographic locality, anatomical site, and host taxonomy, but not host phylogeny

Permutational analysis of variance (PERMANOVA) identified geographic locality, host taxonomy, and anatomical sampling site (gut, oral, skin) as significant factors explaining variation in three independent measures of microbial beta diversity (Bray-Curtis, unweighted UniFrac, and weighted UniFrac) (*p* < 0.001, ADONIS) (Table 3). Measures of intraspecific beta dispersion among weighted UniFrac, unweighted UniFrac, and Bray-Curtis distances showed a continuum of dissimilarities across host species (Fig. S2); mean beta dispersion differed significantly between anatomical sites by all three dissimilarity measures (Dunn’s Test, Bonferroni corrected *p*-value *p* < 0.001).) Analysis of sites by elevation revealed that bats at higher elevations tended to host increased Shannon diversity (SI) and observed richness (OR) across oral (SI: R^2^ = 0.076, *p* < 3.1e-9; OR: R^2^ = 0.038, *p* < 2.5e-5), and skin (SI: R^2^ = 0.16, *p* < 2.2e-16; OR: R^2^ = 0.100, *p* < 2.5e-14) microbiomes (Fig. S3).

**Table 3.** Permutational multivariate analysis of variance using distance matrices, with distance matrices among sources of variation partitioned by host taxonomy (species nested within genus), locality, and anatomical site; * indicates p-value < 0.05.

| Partition Variable | Weighted UniFrac |  |  | Unweighted UniFrac |  |  | Bray-Curtis |  |  |
| --- | --- | --- | --- | --- | --- | --- | --- | --- | --- |
|  | SumSq | F | Pr(>F) | SumSq | F | Pr(>F) | SumSq | F | Pr(>F) |
| Anatomical site | 10.67 | 198.01 | 0.001* | 56.52 | 82.90 | 0.001* | 38.2 | 36.97 | 0.001* |
| Host Genus | 3.77 | 13.09 | 0.001* | 25.54 | 7.02 | 0.001* | 85.30 | 15.06 | 0.001* |
| Locality | 1.56 | 11.00 | 0.001* | 20.62 | 11.34 | 0.001* | 23.85 | 8.42 | 0.001* |
| Host Genus:species | 1.39 | 4.08 | 0.001* | 11.20 | 2.59 | 0.001* | 25.25 | 1.33 | 0.001* |

Across all bat species, the gut microbiome was enriched for Proteobacteria (avg 55.4%) (Enterobacteraceae, avg 50.0%) and Firmicutes (avg 21%) (Clostridiaceae, avg 9.5%; Streptococcaceae, avg 5.5%). Oral microbiota were also enriched for Proteobacteria (avg 64.3%) (Pasteurellaceae, avg 47.5%; Neisseriaceae, avg 8.3%) and Firmicutes (avg 11.4%) (Streptoccaceae, avg 8.8%; Gemellaceae, avg 3.61%). The skin microbiome was not enriched for a single bacterial family, and showed a pronounced increase in relative abundance of Actinobacteria (avg 10%) (Mycobacteraceae, avg 4.1%; Pseudonocardiaceae, avg 2.8%; Nocardiaceae, avg 2.3%) and Bacteroidetes (Moraxellaceae, avg 5.6%), and Euryarchaeota (Halobacteraceae, avg 4.2%) (Fig. 3).

**Figure 3.**
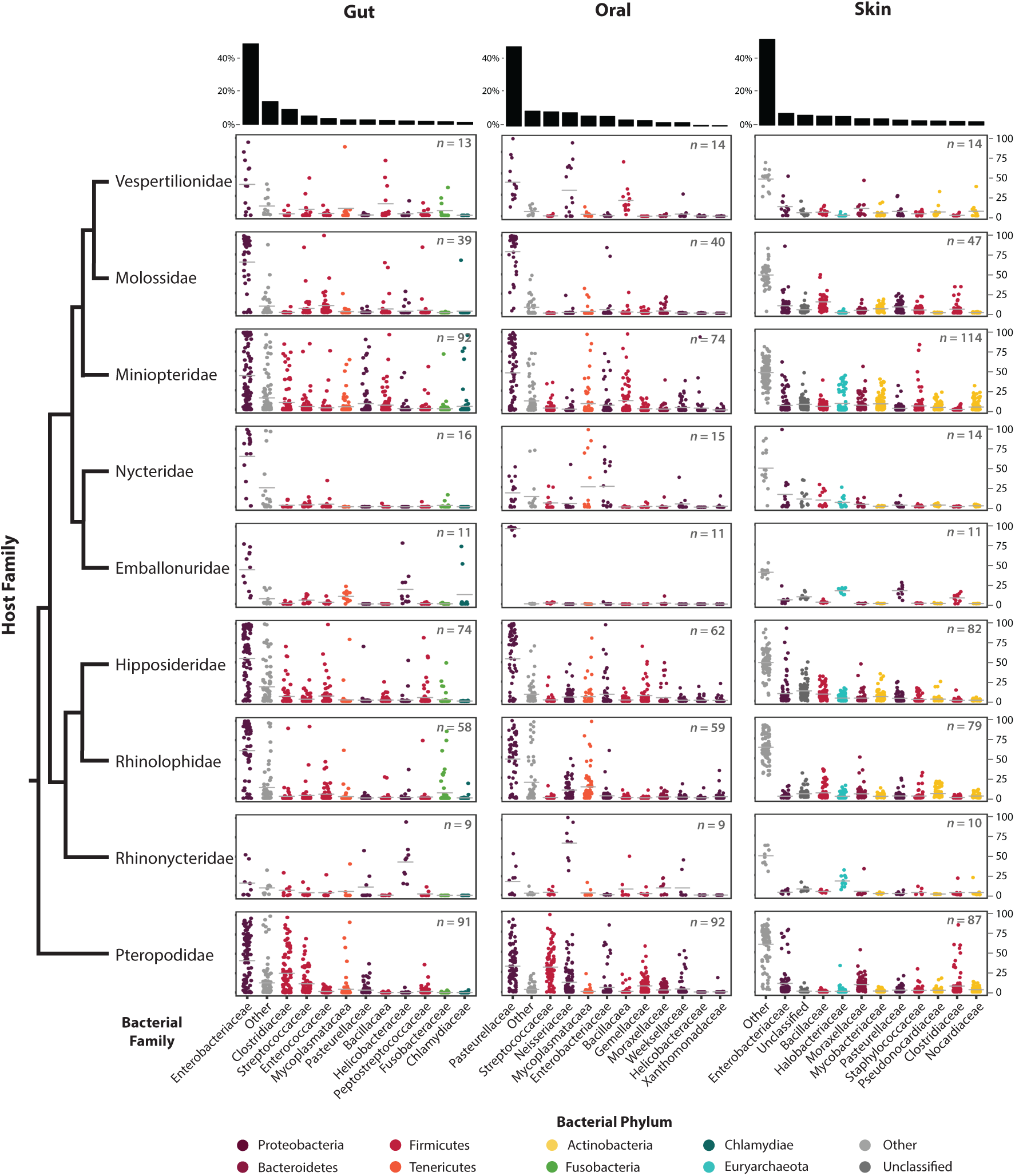
Percent relative abundance of top 11 bacterial families identified in gut, oral, and skin microniomes of bats. Individual points represent the relative abundance of bacterial families within a single library. Results are faceted by anatomical site and arranged by host phylogenetic relationship. Bacterial families are colored according to bacterial phylum. Number of libraries is indicated in the upper right-hand corner of each plot.

Fruit bats (pteropodids) showed enrichment of Clostridiaceae in the gut (avg 24.8%) and Streptococcaeae in the oral microbiome (avg 31.0%) compared to all other bats. The oral microbiota of several insectivorous bat families were enriched for Firmicutes in the Mycoplasmataceae family (nycterids, avg 25.5%; rhinolophids, avg 13.8%; miniopterids, avg 8.4%). The skin microbiota of several insectivorous bat families were enriched for Firmicutes in the Bacillaceae family (molossids, 14.0%; hipposiderids, 8.6%; nycterids, 8.6%; rhinolophids, 6.3%).

Host phylogeny from bat specimens collected during this study was reconstructed to test for significance of phylosymbiosis between bat species and their microbiome (Supplemental Figure 5; Figure 4). Mantel tests of host phylogenetic distances and microbial community dissimilarity (weighted (wuf) and unweighted UniFrac (uf) distances) revealed no correlation for gut (wuf: *R*^2^ = - 0.045, *p* = 0.63; uf: *R*^2^ = - 0.052, *p* = 0.64), oral (wuf: *R*^2^ = - 0.11, *p* = 0.88), and skin (wuf: *R*^2^ = - 0.081, *p* = 0.79; uf: *R*^2^ = - 0.108, *p* = 0.92) microbiota and host phylogenetic distance, with the exception of oral uf dissimilarity and host phylogenetic distance(uf: *R*^2^ = 0.223, *p* = 0.02) (Fig. 4).

**Figure 4.**
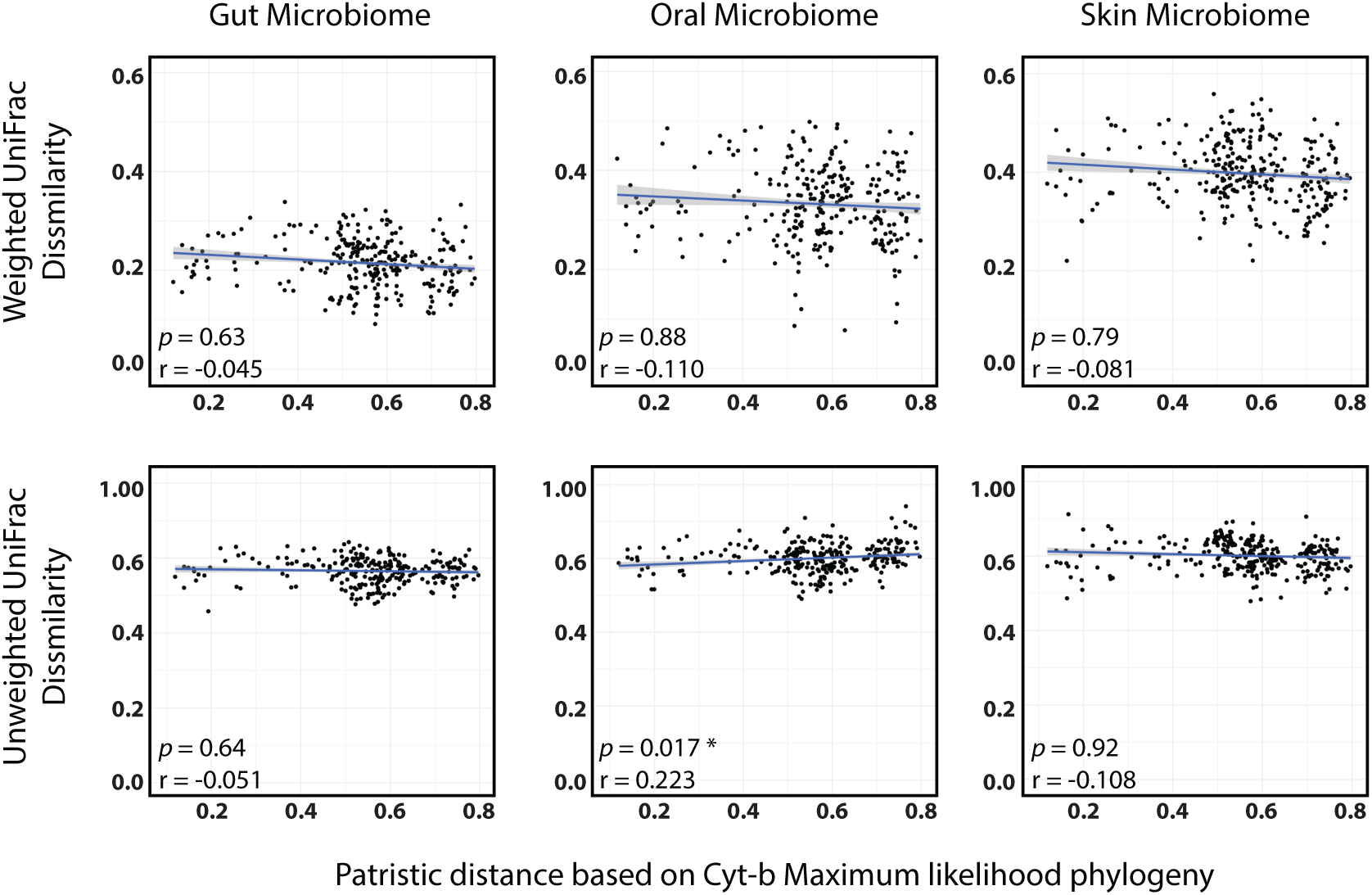
Rate of microbiome divergence across phylogenetic distance of bats. Strengths of correlations assessed by Mantel tests (10,000 permutations) of microbial community dissimilarity (unweighted and weighted UniFrac) and patristic distances calculated from a maximum likelihood hypothesis of bat species from this study. Asterisk indicates significant correlation (p<0.05) as determined by Mantel test.

**Figure 5.**
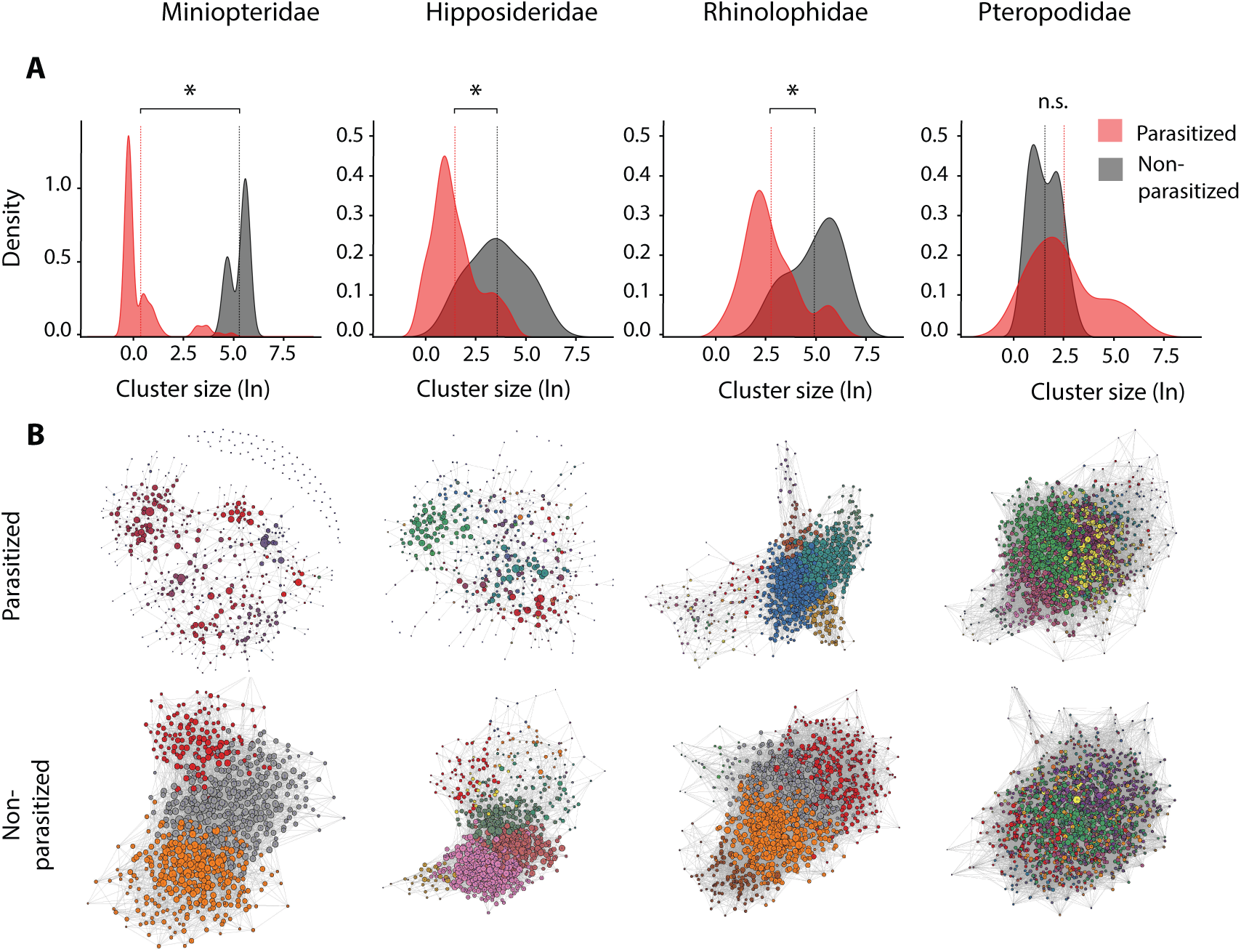
(A) Distribution of skin microbial network clusters for parasitized and non-parasitized bats, grouped by bat family (asterisks indicate signifiance at *p* < 0.005, Kruskal-Wallis) (B) Visualization of skin bacterial networks (based on Fruchterman-Reingold algorithm); colored nodes correspond to unique clusters of co-occurring ASVs within each network.

Patristic distance based on Cyt-b Maximum likelihood phylogeny

### 4 Bat skin microbiome is associated with parasitism in African bats

To test for significant associations between bacterial communities and eukaryotic parasites (obligate ectoparasitic dipteran insects, and obligate endoparasitic malarial parasites), we employed a combination of machine learning techniques, network analyses, and DESeq2 models (see methods). PERMANOVA analysis identified ectoparasite status and malarial infection status as significant predictors of bacterial beta diversity dissimilarity among skin and oral microbiota, respectively (*p* < 0.001, ADONIS). Tests of three independent measures of beta diversity (weighted UniFrac, unweighted UniFrac, and Bray-Curtis) produced congruent results, with the exception of oral microbiome, which was not significantly predictive of malarial infection based on unweighted UniFrac analysis (Table 4).

**Table 4.**
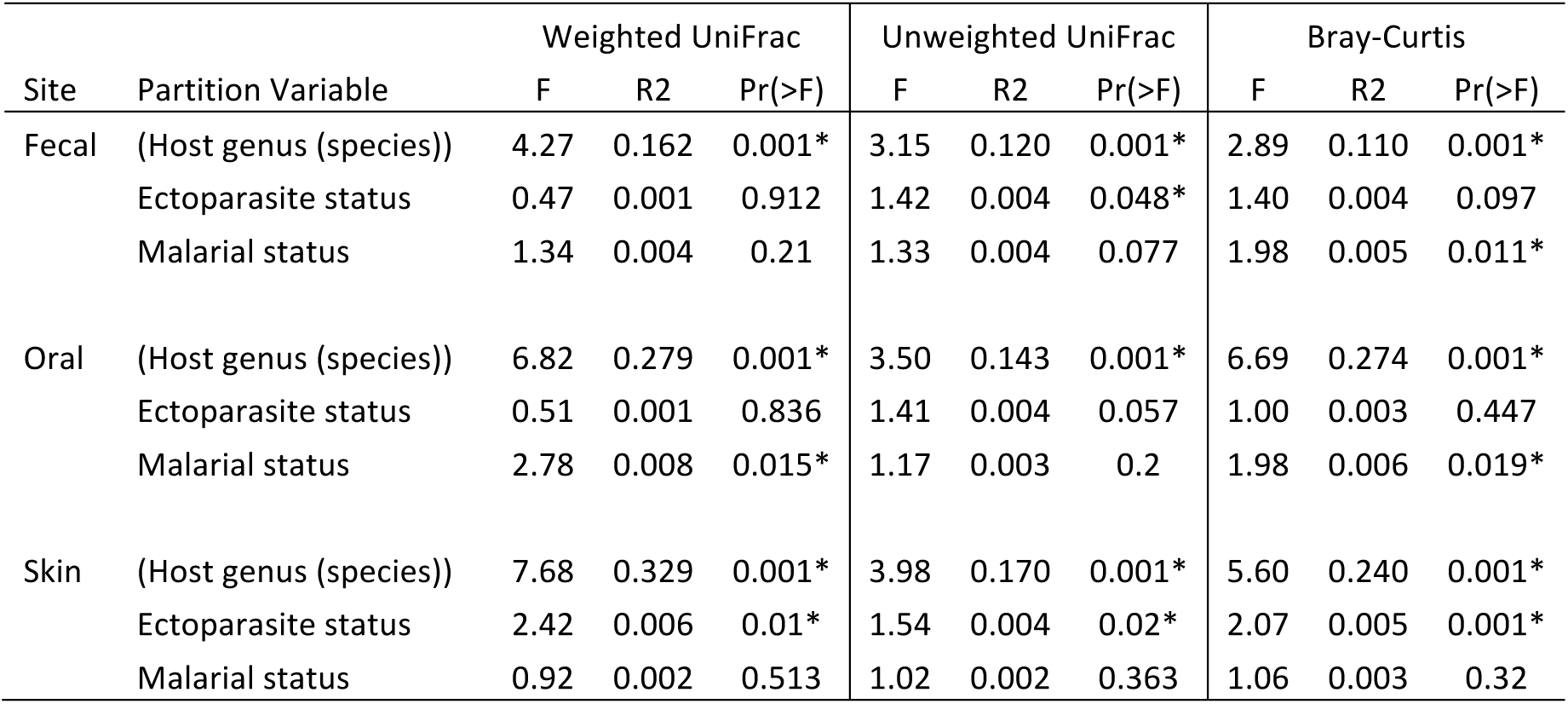
Permutational multivariate analysis of variance using distance matrices, with distance matrices among sources of variation partitioned by host taxonomy (species nested within genus), ectoparasite status, malarial infection status, and locality included as strata; * indicates p-value < 0.05.

| Site | Partition Variable | Weighted UniFrac |  |  | Unweighted UniFrac |  |  | Bray-Curtis |  |  |
| --- | --- | --- | --- | --- | --- | --- | --- | --- | --- | --- |
|  |  | F | R2 | Pr(>F) | F | R2 | Pr(>F) | F | R2 | Pr(>F) |
| Fecal | (Host genus (species)) | 4.27 | 0.162 | 0.001* | 3.15 | 0.120 | 0.001* | 2.89 | 0.110 | 0.001* |
|  | Ectoparasite status | 0.47 | 0.001 | 0.912 | 1.42 | 0.004 | 0.048* | 1.40 | 0.004 | 0.097 |
|  | Malarial status | 1.34 | 0.004 | 0.21 | 1.33 | 0.004 | 0.077 | 1.98 | 0.005 | 0.011* |
| Oral | (Host genus (species)) | 6.82 | 0.279 | 0.001* | 3.50 | 0.143 | 0.001* | 6.69 | 0.274 | 0.001* |
|  | Ectoparasite status | 0.51 | 0.001 | 0.836 | 1.41 | 0.004 | 0.057 | 1.00 | 0.003 | 0.447 |
|  | Malarial status | 2.78 | 0.008 | 0.015* | 1.17 | 0.003 | 0.2 | 1.98 | 0.006 | 0.019* |
| Skin | (Host genus (species)) | 7.68 | 0.329 | 0.001* | 3.98 | 0.170 | 0.001* | 5.60 | 0.240 | 0.001* |
|  | Ectoparasite status | 2.42 | 0.006 | 0.01* | 1.54 | 0.004 | 0.02* | 2.07 | 0.005 | 0.001* |
|  | Malarial status | 0.92 | 0.002 | 0.513 | 1.02 | 0.002 | 0.363 | 1.06 | 0.003 | 0.32 |

Supervised machine learning analyses (random forests) produced models that could classify the anatomical source of microbial communities and the host genus of gut, oral, and skin microbial samples with reasonable accuracy (ratio of baseline to observed classification error ≥2; *i.e.* random forest models performed at least twice as well as random). Random forest models also performed accurately when classifying ectoparasite status based on skin bacterial community composition, but were less accurate for classification of malarial status based on oral bacterial community composition (Table 5).

**Table 5.** Supervised machine learning results, showing random forest model performance with respect to different classification variables and input data sets (fecal, oral, skin microbiome). Model performance is assessed by measuring the ratio of Out-of-bag estimated error (OOB) to baseline error.

| Classification variable | Input Data | Baseline error | OOB error | Baseline:OOB |
| --- | --- | --- | --- | --- |
| Anatomical site | All data | 0.68 | 0.14 | 4.8 |
| Host Genus | Skin | 0.75 | 0.17 | 4.3 |
| Host Genus | Oral | 0.78 | 0.24 | 3.2 |
| Host Genus | Gut | 0.77 | 0.35 | 2.2 |
| Ectoparasite Status | Skin | 0.53 | 0.27 | 2.0 |
| Malarial Status (Miniopteridae only) | Oral | 0.46 | 0.38 | 1.2 |

Following the application of statistical and machine learning approaches, we employed network analyses to characterize the co-occurrence topology of microbial communities (in terms of the relative abundance of co-occurring ASVs) across the skin microbiota of our four most well-sampled bat families (Hipposideridae (*n* = 80), Miniopteridae (*n* = 116), Rhinolophida (*n* = 88), and Pteropodidae (*n* = 106)). Network analyses produced strikingly consistent results, revealing a significant decrease in cluster size (*p* < 0.05, Mann-Whitney-Wilcoxon rank sum test) and median node degree (*p* < 0.05, *t* test), as well as reduced network connectivity for parasitized bats from three of the four bat families examined (Fig. 5; Fig. S4).

### 5 Bacterial taxa on skin correlated with presence or absence of obligate dipteran ectoparasites

DESeq2 analyses of the skin microbiota in four well-sampled bat families (Hipposideridae, Miniopteridae, Rhinolophidae, Pteropodidae) identified a number of ASVs that were significantly associated with either ectoparasitized or non-ectoparasitized bats (Fig. 6). Overall, we identified 89 and 24 ASVs significantly associated with parasitized and non-parasitized bats, respectively (Table S3). Bacterial classes with the greatest representation among significant results were Actinobacteria (16 families), Gammaproteobacteria (11 families), Bacilli (5 families), and Alphaproteobacteria (3 families). ASVs significantly enriched in parasitized bats from at least three out of four bat families included Mycobacteraceae (Actinobacteria), and Xanthomonadaceae (Gammaproteobacteria). ASVs significantly enriched in parasitized bats from at least two out of four bat families included Hyphomicrobiaceae (Alphaproteobacteria), Alcaligenaceae (Betaproteobacteria), Moraxellaceae (Gammaproteobacteria), Planococcaceae (Bacilli), Flavobacteraceae (Flavobacteria), Halobacteraceae (Halobacteria), and Chitinophagaceae (Saprospirae) (Fig. 6).

**Figure 6.**
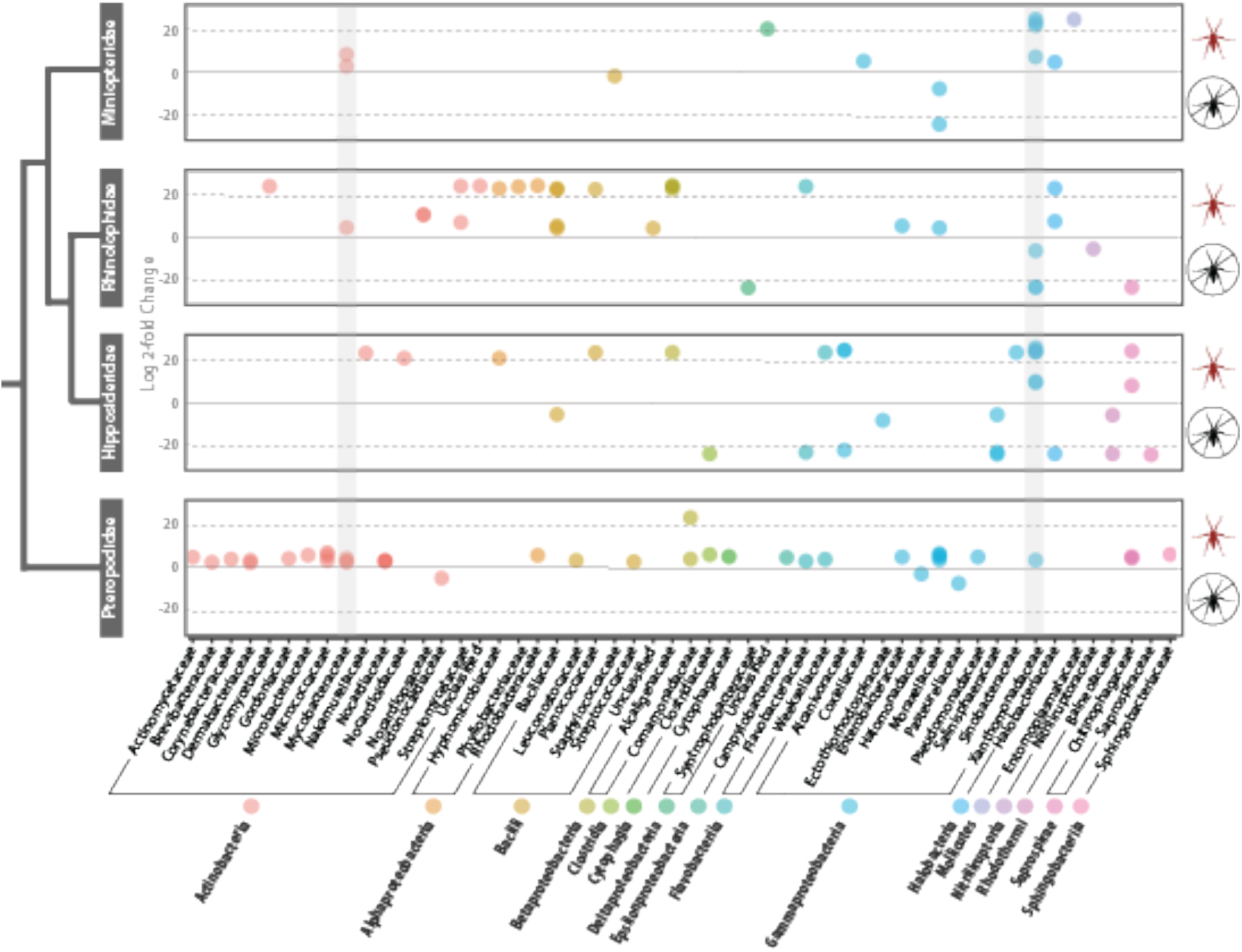
Log2fold change in relative abundance of skin-associated ASVs from the four most-sampled bat families estimated with DESeq2. ASVs shown were found to be significantly associated with ectoparasite status in bats based on analysis of negative binomial distributions of relative abundance (Banjamini-Hochberg FDR corrected *p*-value *p* < 0.05). Positive values correspond to ASVs found to be enriched on parasitized bats, and negative values correspond to ASVs found to be enriched on non-parasitized bats. Gray bars highlight ASVs in bacterial families that were enriched in parasitized bats for three out of four bat families.

## DISCUSSION

The bacterial diversity we observed among gut, oral, and skin microbiota of bats fall within ranges similarly observed in other vertebrate groups (3, 20-23). Although few studies have simultaneously compared gut, oral, and skin microbiota from the same individuals, our data reflect an apparent trend in the literature of skin bacterial diversity among vertebrates significantly outnumbering gut or oral bacterial diversity (24-27). Our data corroborate the findings of Nishida and Ochman (3), revealing no relationship between chiropteran phylogeny and gut bacterial community dissimilarity. We also found the same absence of phylogenetic signal among oral and skin microbial communities. As suggested in other studies of volant vertebrates (bats and birds), convergent adaptations driven by the evolution of flight may be influencing the nature and composition of microbial communities in both bats and birds (28-30). This differs markedly from studies of other non-volant mammals, such as primates and rodents, for which phylogenetic relatedness is generally a significant predictor of microbial community dissimilarity (21, 31-33).

Microbial community specificity can be assessed as a function of intraspecific variation in dissimilarity (beta dispersion), where a low variance of dispersion suggests a tight and perhaps co-evolutionary link between hosts and symbionts, and a high variance of dispersion suggests more random or non-specific associations between hosts and symbionts (34). Measures of beta dispersion among bat species revealed a continuum for all three anatomical sites, with oral bacterial communities showing lower levels of beta dispersion than gut or skin communities (Fig. S2). Given that we found no correlation between bacterial community dissimilarity and host phylogenetic distance, and that we observed no taxonomic clustering of hosts in mean beta dispersion estimates, variation in beta dispersion is likely driven by ecological rather than evolutionary factors.

Similar to recent studies in North American bats (35), we found sampling locality to be a significant factor influencing skin, gut and oral microbial composition (Table 3). Furthermore, we observed an apparent trend in increasing Shannon diversity and observed ASV richness along an elevational gradient that was most pronounced for skin microbiota (Fig. S3). A positive correlation between bacterial richness and elevation has been observed in studies of amphibian skin (36) and montane soil, and this pattern may be the result of climatological and other abiotic factors (*e.g.* pH) found along elevational gradients (37, 38).

We found the general composition of gut microbiota in East African bats to be similar to that of Neotropical bats, with Proteobacteria being the dominant bacterial phylum present (39). Regardless of diet (insectivorous or frugivorous), the distal bat gut is dominated by bacteria in the family Enterobacteriaceae (Phylum: Proteobacteria), though fruit bats do have an increased relative abundance of bacteria in the family Clostridiaceae (Phylum: Firmicutes) relative to insectivorous bats. In their study of neotropical bats, Phillips et al. (40) noted an increased relative abundance of Lactobacillales in frugivorous bats, and we note a similar pattern among pteropodid fruit bats in this study, which exhibited a slightly higher proportion of Streptococcaceae (Order: Lactobacillales) relative to insectivorous bats (Fig. 3). Overall, the predominant enrichment of the chiropteran gut by Proteobacteria differs markedly from other mammalian gut microbiomes, which are generally dominated by Firmicutes (21, 41, 42).

Among most bat families, the oral microbiome was dominated by Pasteurellaceae (Phylum: Proteobacteria), and in some cases a high relative abundance of bacteria in the families Mycoplasmataceae (in nycterids), Neisseriaceae (in vespertilionids and rhinonycterids), and Streptococcaceae (in pteropodids) was also observed. Although the oral microbiome has received less attention than that of the gut, several studies have found diverse Pasteurellaceae and Neiserria lineages present in the oral microbiota of animals, including domestic cats (20) and marine mammals (43). Pasteurellaceae lineages have also recently been documented in the oral microbiota of Tasmanian devils (23, 44). In humans, Pasteurallaceae (genera *Haemophilus* and *Aggregatibacter*) and Neisseriaceae (genera *Neisseria*, *Kingella*, and *Eikenella*) play an important role in the formation supragingival plaque (22). Though these bacterial groups are present in lower proportions in other animals relative to bats, their presence in a broad range of host taxa suggest a conserved evolutionary niche.

Our analysis identified links between ectoparasitism, malarial parasitism, and bacterial communities on the skin and in oral cavities, respectively. Network analyses identified consistent, stable, and species-rich clusters of bacteria on the skin of non-ectoparasitized bats, compared to relatively disconnected and apparently transient bacteria on the skin of bats harboring ectoparasites. This result mirrors that found in human-mosquito interactions, in which individuals with lower bacterial diversity on the skin are significantly more attractive to blood-seeking mosquitoes than individuals with higher diversity (45). In humans, skin bacteria play a known role in attracting mosquitoes via their production of volatile organic compounds (VOCs), and studies have shown that variation in skin microbial community composition can increase or decrease human attractiveness to blood-seeking mosquitoes (45-47). Similar mechanisms may be at play in the bat-ectoparasite system, particularly given the phylogenetic proximity of hippobscoid bat parasites to mosquitoes.

Several bacterial families exhibited significant associations with presence of ectoparasitism in bats based on DESeq analyses. Bacteria found across multiple host families included (but were not limited to) Alcaligenaceae, Chitinophagaceae, Flavobacteriaceae, Moraxellaceae, Mycobacteriaceae (*Mycobacterium* spp.), and Xanthomonadaceae. In many cases, these bacterial families were associated with parasitism in some bat families, and absence of parasitism in others, suggesting a potential mechanism by which ectoparasites might be distinguishing between “correct” and “incorrect” hosts. As suggested by human-mosquito interaction studies (45, 46, 48), bacteria positively associated with increased rates of blood-feeding dipteran host selection may be producing VOCs on which the insects rely to identify their hosts. Bacteria that are negatively associated with such insects may be consuming the products of the former, or may be producing VOCs of their own that mask those of the former (suggested by Verhulst et al. (45)). To better understand the mechanisms underlying these correlations in wild populations, future experiments should consider including sampling of VOCs *in vivo*.

PERMANOVA analyses identified associations between the oral microbiome and malarial parasite prevalence among bats in the family Miniopteridae, although these associations were less robust than those of the skin bacteria and ectoparasitism. Upon further exploration of this potential association, we identified a single bacterial ASV in the genus *Actinobacillus* (99% similar to *A. porcinus* based on NCBI blastn search) as significantly reduced in malaria-free bats (baseMean 7.61, −24.2 log2FoldChange, *p* = 1.7E-20). Network analyses indicated no significant differences in connectivity or node degree distribution (results not shown). Because no other bat groups experienced rates of malarial parasitism adequate for statistical analyses, we were unable to explore this relationship further. Future studies that incorporate greater sampling of malaria-positive species may reveal more robust microbial associations, as have been documented in numerous experiments with controlled rodent and human malaria infections (48-50).

Although we cannot ascertain causality of differences in the microbial composition of skin in this study, our results support the hypothesis that these differences suggest a mechanism by which ectoparasites can locate or distinguish hosts. Alternatively, observed differences in microbial composition could result from microbial transfer from parasites to hosts. Given the known effect of locality and apparent absence of host phylogenetic signal in microbial community composition of skin, one possible explanation is that local environmental variables play a greater role in determining host-bacteria associations in bats. Indeed, in North America, multiple bat species have been found to share many bacterial genera with soil and plant material (35). Thus, local conditions and bacterial composition of bat roosts are likely playing an important role in driving the composition of skin bacteria of bats.

## METHODS

### 1 Sampling

Sampling for this study was conducted from the eastern coast of Kenya to the northern border of Uganda during August-October 2016 (Fig. 1; Table S1, S2). Nine families and nineteen genera of bats (order: Chiroptera) were collected as part of bird and small mammal biodiversity inventories. All sampling was conducted in accordiance with the Field Museum of Natural History IACUC and voucher specimens are accessioned at the Field Museum of Natural History (Table S2). Blood samples were collected and screened for haemosporidia, and haemosporidian taxonomy was assigned using previously described molecular methods (13). Following blood sampling, ectoparasites were removed with forceps and placed directly into 95% EtOH; ectoparasites taxonomy was assigned based on morphological features. For the purposes of analysis with microbiome data, ectoparasite and malarial status were each scored separately as 1 (present) or 0 (absent). Gut, skin, and oral samples were taken for each bat for microbial analyses. Gut samples consisted of fecal material collected directly from the distal end of the colon using sterilized tools, and preserved on Whatman® FTA® cards for microbiome analyses. For oral microbiome analyses, we preserved both buccal swabs in lN_2_ and tongue biopsies in 95% ethanol (EtOH). Comparison of ASV diversity obtained from paired subsets of each sample type revealed greater diversity recovered from tongue biopsies (data not shown); tongues were therefore used for characterization of oral microbiomes in this study. Lastly, skin samples from five regions of the body (ear, wing membrane, tail membrane, chest, back) were collected and pooled in 95% EtOH using sterile Integra® Miltex® 5mm biopsy punches. The goal of sampling from five body regions was to maximize bacterial diversity recovered from the external skin surface of each individual. We based our storage media selections on the recent study by Song et al. (51).

### 2 Microbiome sequencing, characterization, and parasite association

DNA extractions were performed on gut, tongue, and skin samples using the MoBio PowerSoil 96 Well Soil DNA Isolation Kit (Catalog No. 12955-4, MoBio, Carlsbad, CA, USA). We used the standard 515f and 806r primers (52-54) to amplify the V4 region of the 16S rRNA gene, using mitochondrial blockers to reduce amplification of host mitochondrial DNA. Sequencing was performed using paired-end 150 base reads on an Illumina HiSeq sequencing platform. Following standard demultiplexing and quality filtering using the Quantative Insights Into Microbial Ecology pipeline (QIIME2) (55) and vsearch8.1 (56), ASVs were identified using the Deblur method (19) and taxonomy was assigned using the Greengenes Database (May 2013 release; http://greengenes.lbl.gov). According to a recent stuy by McMurdie and Holmes (57), rarefying 16s data is inappropriate for the detection of differentially abundant species. However, for the purposes of comparison, we compared both libraries rarefied to a read depth of 10,000 reads and libraries filtered to those containined >1000 reads (negative controls all contained fewer than 1000 reads and were filtered at this step). Alpha and beta-diversity analyses produced statistically similar results, with no significant differences observed betweeh the rarefied and non-rarefied data. We thus chose to report results of non-rarefied data, based on these observations and the recommendation of McMurdie and Holmes (57).. Following filtering, data were subset for analyses according to sample type, host genus, and locality (or some combination thereof). Site-specific analyses were only performed for sites from which five or more individual bats were sampled. We calculated alpha diversity for each sample type (gut, oral, skin) using the Shannon index, and measured species richness based on actual observed diversity. Significance of differing mean values for each diversity calculation was determined using the Kruskal-Wallis rank sum test, followed by a post-hoc Dunn test with bonferroni corrected *p*-values. Three measures of beta diversity (unweighted UniFrac, weighted UniFrac, and Bray-Curtis) were calculated using relative abundances of each ASV (calculated as ASV read depth over total read depth per library). Significant drivers of communitity similarity were identified using the ADONIS test with Bonferroni correction for multiple comparisons using the R package Phyloseq (58). To assess potential effect of imbalanced sampling, the ADONIS test was re-run on a subset of data comprising only data from the top four sampled bat families, which represented even sampling among families and across the localities from which they were collected. Results of this test (not reported) indicated the same significant drivers of community similarity as the test run on the entire data set. Additional R packages used for analyses and figure generation included vegan (59), ggplot2(60), and dplyr(61). For a complete list of packages and code for microbiome analyses, see http://github.com/hollylutz/BatMP.

### 3 Bat phylogenetic reconstruction

DNA from bats collected during this study was extracted and sequenced for mitochondrial Cytochrome-*b* (cyt-*b*), using the primer pair LGL 765F and LGL 766R that amplify the entire cyt-*b* gene (Bickham et al. 1995, 2004). DNA extractions, PCR amplification, and sequencing were carried out as in Demos et al. 2018. The best-supported model of nucleotide substitution for cyt-*b* was determined using the BIC on the maximum-likelihood topology estimated independently for each model in jMODELTEST2 (Darriba et al., 2012) on CIPRES Science Gateway v.3.1 (Miller et al., 2010). Maximum-likelihood estimates of cyt-*b* gene trees were made using the program IQ-TREE version 1.6.0 (Nguyen et al. 2017) on the CIPRES portal. Emballonuridae (*Coleura afra*) was constrained as sister to Nycteridae (*Nycteris arge*, *N. thebaica*; Amador et al. 2016). We conducted analyses using the ultrafast bootstrap algorithm and searched for best-scoring ML tree algorithm under the GTR+I+ FreeRate model with 1000 bootstrap replicates. The resulting phylogenetic hypothesis and node support can be viewed in Fig. S5.

### 4 Machine learning and network analyses

A supervised machine learning approach was used to produce random forests (RF) for the classification of different variables. RFs were constructed using 1000 decision trees and subsets of ASV data via the supervised_learning.py script implemented in QIIME (55), which utilized 80% of each input data set to train classification models, and tested the accuracy of the models on the remaining 20% of data. We tested the ability of RFs to accurately classify 1) anatomical site (using all data), 2) host genus (using gut, oral, or skin microbial data separately), 3) ecotparasite status (using skin microbial data), and 4) malarial status (using oral microbial data). Classification categories comprised approximately equal numbers of samples, with the exception of host genera, which varied substantially (see Table 1). RF performance was assessed by comparing the out-of-bag estimated error (OOB) with baseline (random) error. If the ratio of OOB to baseline error was less than or equal to two, the model was considered to perform reasonably well, as it performed at least twice as well as random (62). To reconstruct microbial networks for skin and oral bacterial communities within bat family groupings (which were further sub-divided into parasitized or non-parasitized), we utilized the R package Sparse Inverse Covariance Estimation for Ecological Association Inference (SPIEC-EASI) (63). All network datasets were filtered to contain only ASVs that appeared in at least three individuals within each respective dataset. We used the neighborhood selection framework (MB method) with 20 repetitions. Network results produced with SPIEC-EASI were summarized using the R packages CAVnet (64) and igraph (65). Network stability was assessed by sequentially removing network nodes (ordered by betweeness centrality and degree) and observing natural connectivity (*i.e.* eigenvalue of the graph adjacency matrix) as nodes are removed. To determine which, if any, bacterial ASVs were significantly associated with ectoparasite or malarial prevalence, we performed analyses based on the negative binomial distribution of ASVs relative abundance, utilizing the R package DESeq2 (66). For ectoparasite-assocation tests, the data were subset into four categories that corresponded to the top-sampled bat families (Hipposideridae, Miniopteridae, Pteropodidae, and Rhinolophidae), each with similar propotions of parasitized to non-parasitzed individuals (see Table 1). For haemosporidian-associated tests, only the family Miniopteridae was analyzed, due to highly imbalanced prevalence or sample sizes across all other families (Table 1). Dispersion estimates and fit tests were implented using default DESeq2 parameters. False discovery rate (FDR) was calculated using the Benjamini-Hochberg method for each of the bat families analyzed, and *p*-values were adjusted accordingly.

## ACKNOWLEDGMENTS

We thank the Kenya Wildlife Service and the Uganda Wildlife Authority for permission to conduct research in national parks. For logistical support and assistance in the field, we thank Mike Bartonjo of the National Museums of Kenya, Phausia Kweyu of Karatina University, Dr. Robert Kityo, Solomon Sebuliba, and Cissy Akoth of the Makerere University Zoological Museum, Drs. Brian Amman, Jonathan Towner, and Rebecca Tiller of the Centers for Disease Control and Prevention, and Lauren Lutz. We thank Neil Gottel for his knowledge and assistance with laboratory processing of microbial samples, and other members of the Gilbert Lab, including Alyson Yee, Cesar Cardona, Thomas Kuntz, Drs. Bea Penalver, Melissa Dsouza, and Naseer Sangwan for the assistance with bacterial 16s analyses.

## AUTHOR CONTRIBUTIONS

H.L.L. designed the research and wrote the first draft; H.L.L., E.W.J. analyzed data; T.C.D. produced bat phylogeny; H.L.L., P.W.W., W.B.S., J.C.K. conducted field research; J.A.G., B.D.P. provided funding and research support; all authors contributed to writing.

## Supplemental Figures

**Figure S1.**
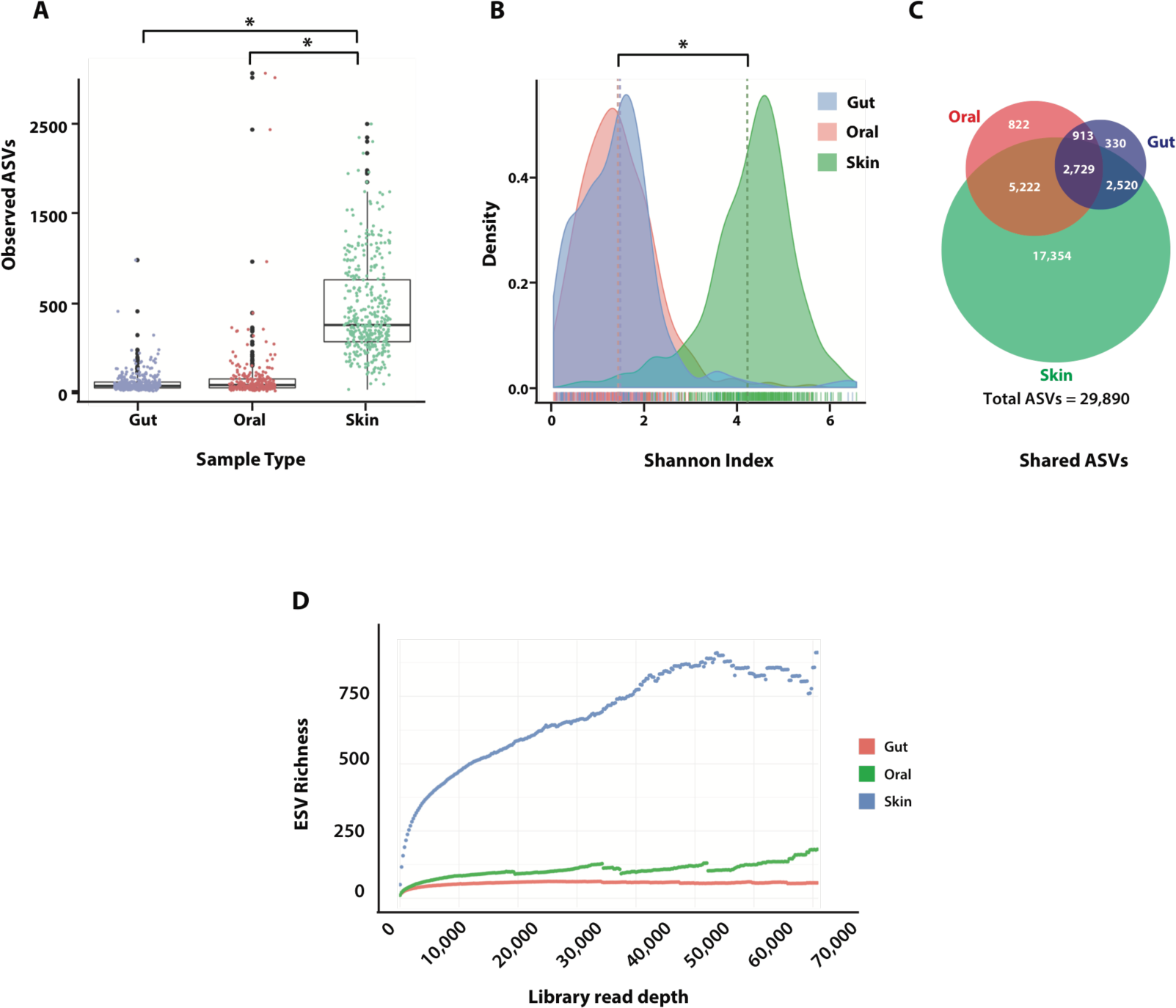
Alpha diversity measures of data rarefied to a read depth of 10,000. A) Observed ASV richness across anatomical site; gut and oral microbial richness differeed significantly from skin microbial richness (Kruskal-Wallis chi-squared = 677.01, df = 2, p < 2.2e-16, Dunn’s test), but did not differ significantly from each other. B) Density plots of Shannon Index (SI) by anatomical site; SI of skin microbial richness and evenness differed significantly from the SI of both gut and oral microbiomes, which did not differ from each other (Kruskal-Wallis chi-squared = 678.0885, df = 2, p < 2.2e-16, Dunn’s test). C) Venn diagram of shared and specific ESVs across different anatomical sites. As with analyses of the non-rarefied data set, skin exhibited the high-est diversity (27,825 ASVs), followed by the oral microbiome (10,696 ASVs), and lastly the gut (6,492 ESVs). Rarefying data led to a loss of 1,079 ASVs (4% of total ASVs) that did not appear in any sample after rarefaction, and the removal of 1,079 libraries that had <10,000 reads. D) Mean ASV richness as read depth increases, with removal of libraries containing fewer than the identified number of reads (note - color key in D differs from color key in A-C).

**Figure S2.**
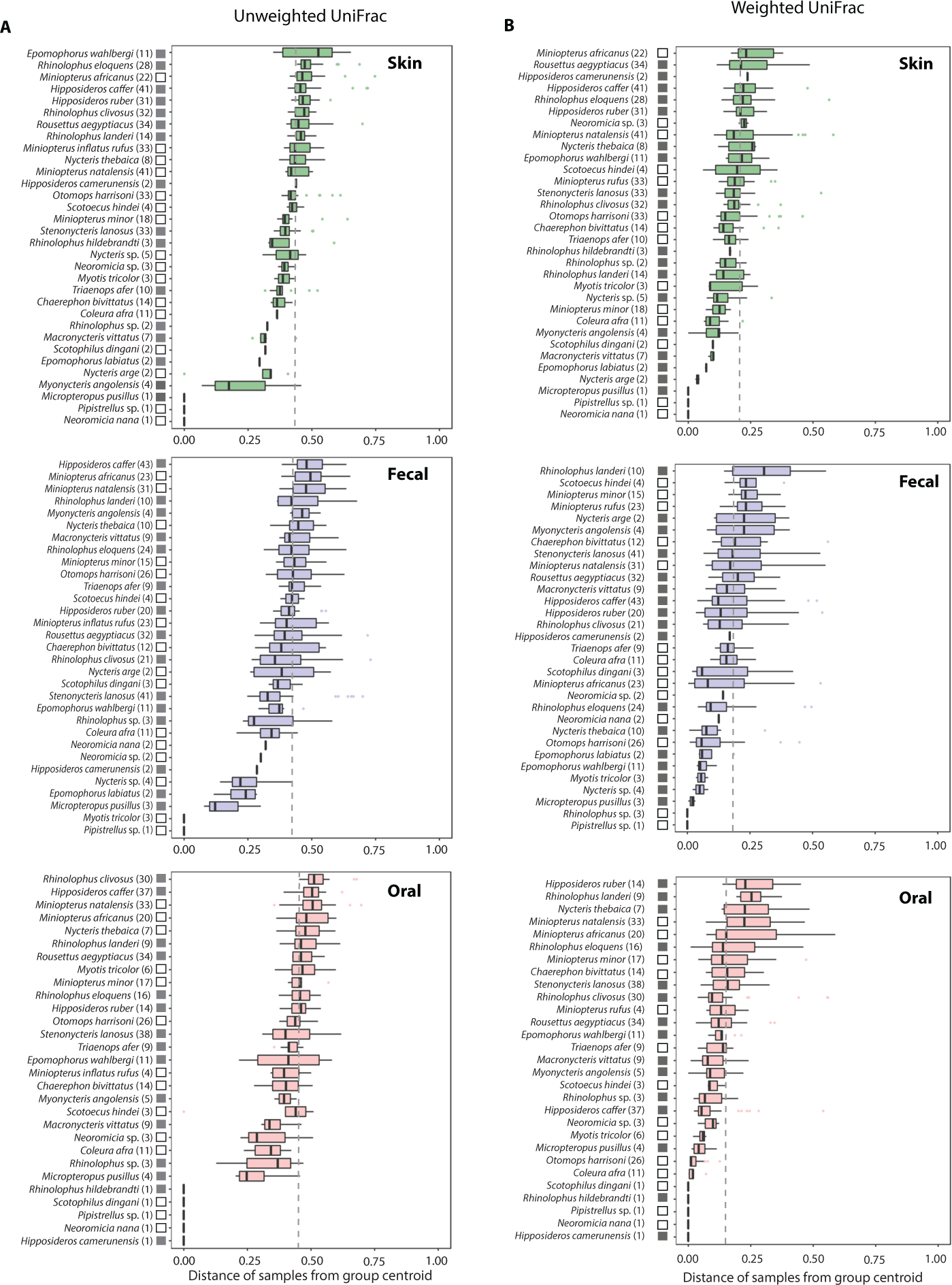
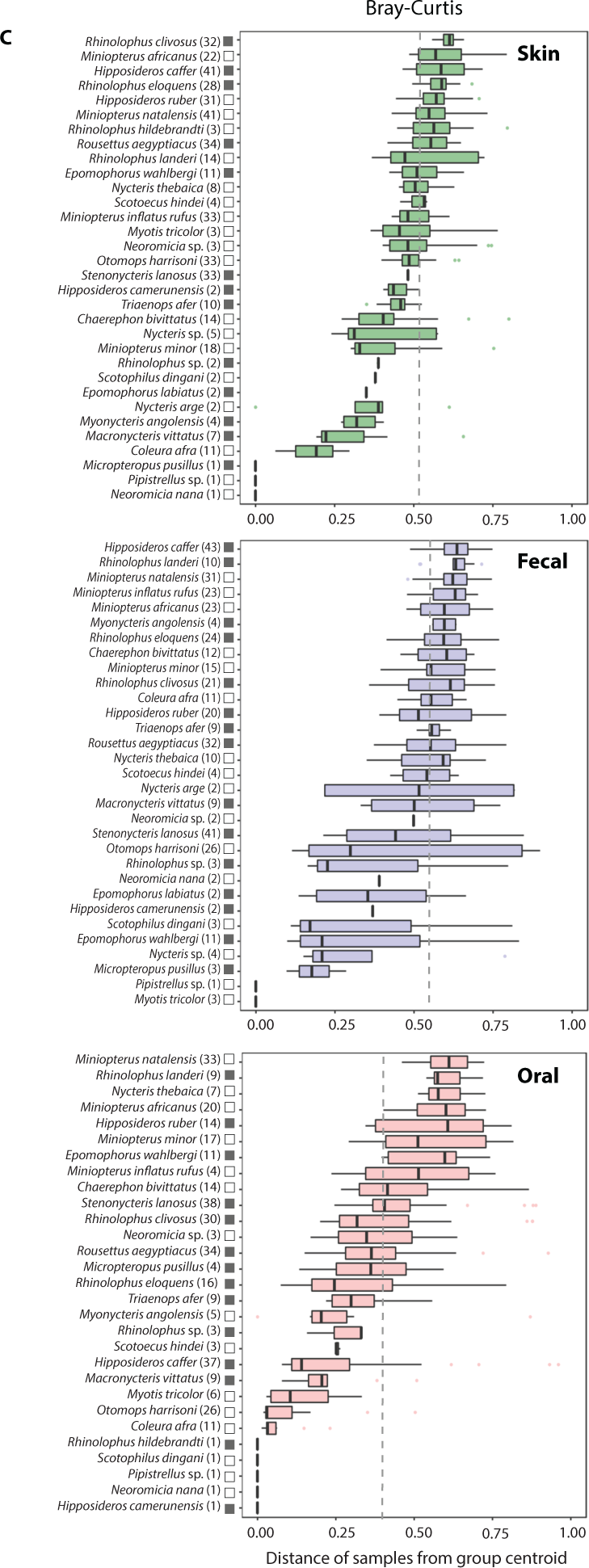
Intraspecific variation across anatomical sites measured as beta dispersion of (A) unweighted UniFrac, (B) weighted UniFrac, and (C) Bray-Curtis distances. Dotted lines indicate mean dispersion for anatomical groupings; numbers in parentheses indicate sample size per bat species. White and gray boxes correspond to the chiropteran suborders Yangochiroptera (microbats) and Yinpterochiroptera (fruits bats and kin), respectively.

**Figure S3.**
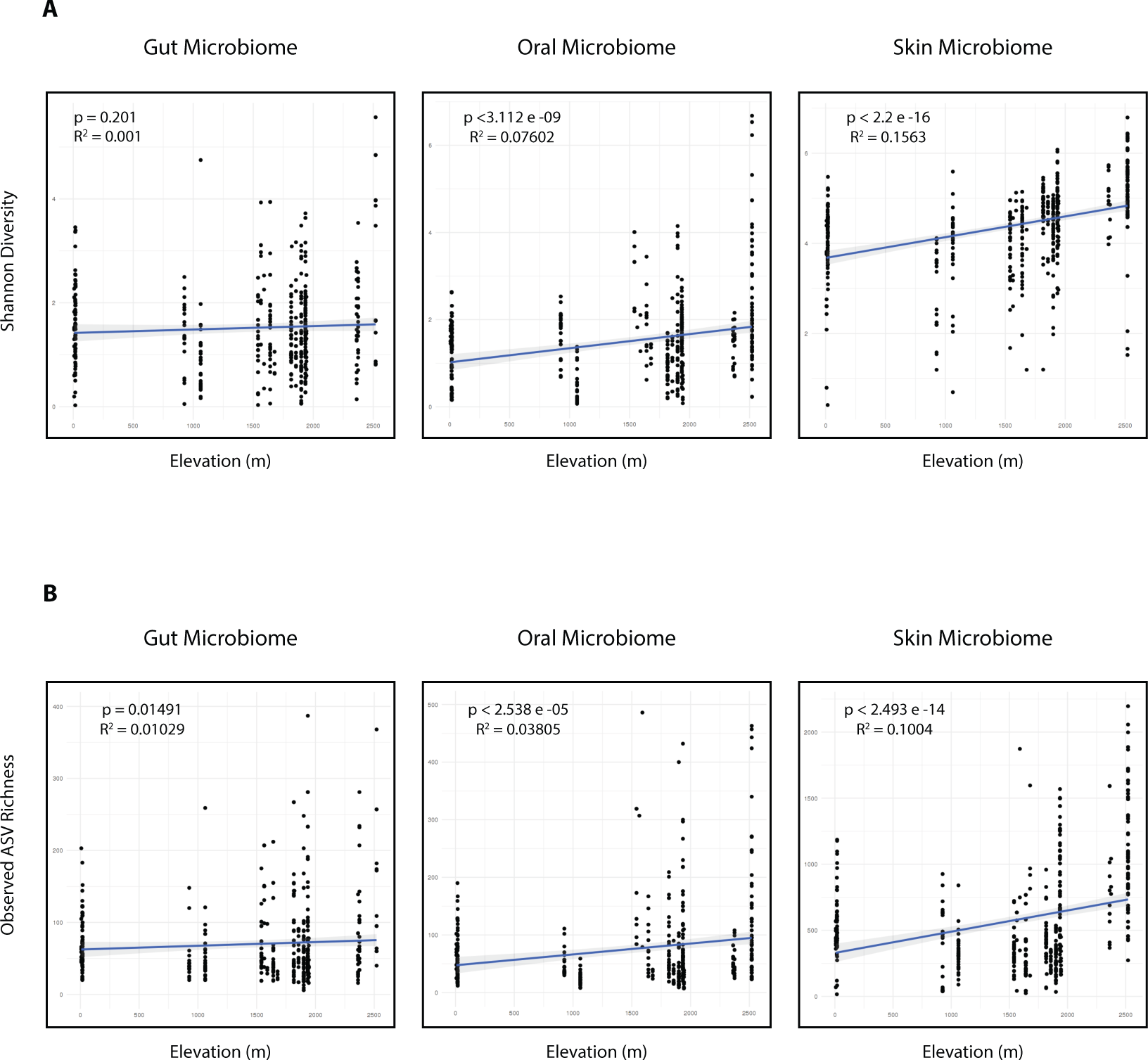
Linear regression of (A) Shannon diveristy and (B) observed “SV richness of gut, oral, and skin microbiomes against elevation from which host was sampled (∼0 - 2500 meters above sea-level). R^2^ and significance values are provided within each plot.

**Figure S4.**
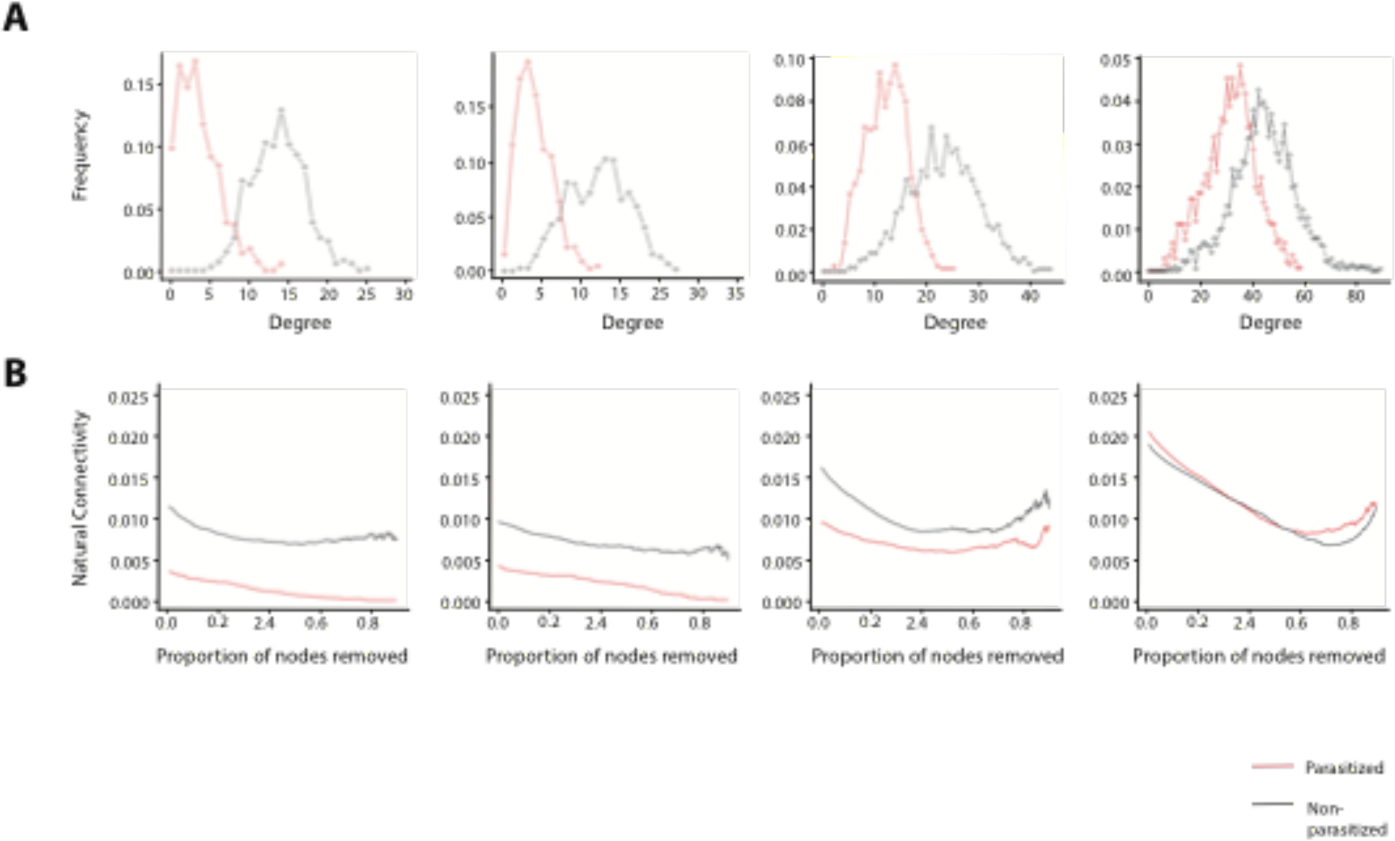
Network analyses. A) Node degree distribution of parasitized and non-parasitized bats,grouped by family.B) Network fragility plots,showing natural network connectivity with sequential removal of nodes ordered by betweenness and degree.

**Figure S5.**
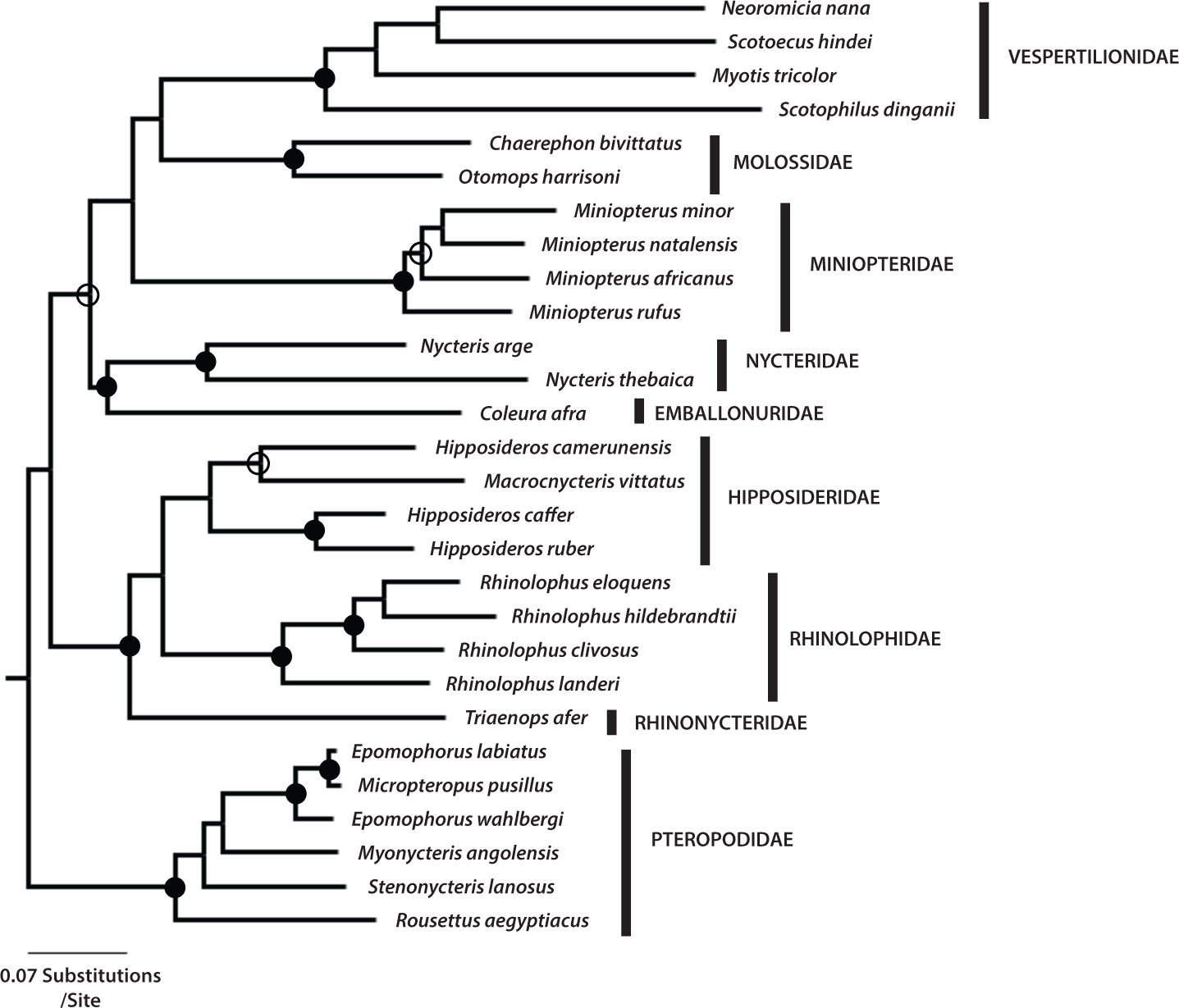
Maximum likelihood phylogeny of bat species Cased on the mitochondrial Cytochrome *b* locus (Cyt *b*). Phylogenetic distances were calculated as patristic distances based on maximum likelihood reconstruction of bat species-level phylogeny with 1000 bootstrap (bs) replicates. Closed CMBDL circles > 97% bs support, open circles > 70% bs support. Voucher specimens are accessioned at the Field Museum of Natural History (Chicago, IL); accession information can be found in Table S3 (where specimens included in phylogenetic analyses BSF highlighted in red).

## REFERENCES

1. Human Microbiome Project C (2012) Structure, function and diversity of the healthy human microbiome. Nature 486(7402):207–214.

2. Thaiss CA, Zmora N, Levy M, & Elinav E (2016) The microbiome and innate immunity. Nature 535(7610):65–74.

3. Nishida AH & Ochman H (2018) Rates of gut microbiome divergence in mammals. Mol Ecol 27(8):1884–1897.

4. Moeller AH, et al. (2014) Rapid changes in the gut microbiome during human evolution. Proc Natl Acad Sci U S A 111(46):16431–16435.

5. Kundu P, Blacher E, Elinav E, & Pettersson S (2017) Our Gut Microbiome: The Evolving Inner Self. Cell 171(7):1481–1493.

6. Li X, et al. (2017) Composition of Gut Microbiota in the Gibel Carp (Carassius auratus gibelio) Varies with Host Development. Microb Ecol 74(1):239–249.

7. Kolodny o, et al. (2017).

8. McFall-Ngai M, et al. (2013) Animals in a bacterial world, a new imperative for the life sciences. Proc Natl Acad Sci U S A 110(9):3229–3236.

9. Kwiatkowski DP (2005) How malaria has affected the human genome and what human genetics can teach us about malaria. Am. J. Hum. Genet. 77:171– 190.

10. Dittmar K, Morse, Solo F., Dick, Carl W., Patterson, Bruce D. (2015) Bat fly evolution from the Eocene to the present (Hippoboscoidea, Streblidae and Nycteribiidae). Parasite Diversity and Diversification: Evolutionary Ecology Meets Phylogenetics, ed S. Morand BRK, D. T. J. Littlewood (Cambridge University Press, Cambridge, U. K.).

11. Dick CW, Patterson, B. D. (2006) Micromammals and Macroparasites: From Evolutionary Ecology to Management (Springer, Kato Bunmeisha, Japan).

12. Schaer J, et al. (2013) High diversity of West African bat malaria parasites and a tight link with rodent Plasmodium taxa. Proceedings of the National Academy of Sciences 100:17415–17419.

13. Lutz HL, et al. (2016) Diverse sampling of East African haemosporidians reveals chiropteran origin of malaria parasites in primates and rodents. Mol Phylogenet Evol 99:7–15.

14. owner JS, et al. (2009) Isolation of genetically diverse Marburg viruses from Egyptian fruit bats. PLoS Pathog 5(7):e1000536.

15. Olival KJ & Hayman DT (2014) Filoviruses in bats: current knowledge and future directions. Viruses 6(4):1759–1788.

16. Amman BR, et al. (2015) A Recently Discovered Pathogenic Paramyxovirus, Sosuga Virus, is Present in Rousettus aegyptiacus Fruit Bats at Multiple Locations in Uganda. J Wildl Dis 51(3):774–779.

17. Li W, et al. (2005) Bats are natural reservoirs of SARS-like coronaviruses. Science 310:676–679.

18. Chua KB, et al. (2002) Isolation of Nipah virus from Malaysian island flying-foxes. Microbes and Infection 4:145–151.

19. Amir A, et al. (2017) Deblur Rapidly Resolves Single-Nucleotide Community Sequence Patterns. mSystems 2(2).

20. Sturgeon A, Pinder SL, Costa MC, & Weese JS (2014) Characterization of the oral microbiota of healthy cats using next-generation sequencing. Vet J 201(2):223–229.

21. Ley RE, et al. (2008) Evolution of mammals and their gut microbes. Science 320(5883):1647–1651.

22. Mark Welch JL, Rossetti BJ, Rieken CW, Dewhirst FE, & Borisy GG (2016) Biogeography of a human oral microbiome at the micron scale. Proc Natl Acad Sci U S A 113(6):E791–800.

23. Brix L, Hansen MJ, Kelly A, Bertelsen MF, & Bojesen AM (2015) Occurrence of Pasteurellaceae Bacteria in the Oral Cavity of the Tasmanian Devil (Sarcophilus Harrisii). J Zoo Wildl Med 46(2):241–245.

24. Grice EA & Segre JA (2011) The skin microbiome. Nat Rev Microbiol 9(4):244– 253.

25. Ursell LK, et al. (2012) The interpersonal and intrapersonal diversity of human-associated microbiota in key body sites. J Allergy Clin Immunol 129(5):1204–1208.

26. Costello EK, et al. (2009) Bacterial community variation in human body habitats across space and time. Science 326(5960):1694–1697.

27. Chiarello M, Villéger, S., Bouvier, C., Bettarel, Y., Bouvier, T. (2015) High diversity of skin-associated bacterial communities in marine fishes is promoted by their high variability among body parts, individuals and species. FEMS Microbiol Ecol 91(7):1–12.

28. Caviedes-Vidal E, McWhorter, T. J., Lavin, S. R., Chediak, J. G., Tracy, C. R., Karasov, W. H. (2007) The digestive adaptation of flying vertebrates: High intestinal paracellular absorption compensates for smaller guts. PNAS 104(48):19132–19137.

29. Price ER, Brun A, Caviedes-Vidal E, & Karasov WH (2015) Digestive adaptations of aerial lifestyles. Physiology (Bethesda) 30(1):69–78.

30. Caviedes-Vidal E, et al. (2008) Paracellular absorption: a bat breaks the mammal paradigm. PLoS One 3(1):e1425.

31. Moeller AH, Car-Qintero, A., Mjungu, D., Georgiev, A. V., Lonsdorf, E. V., Muller, M. N., Pusey, A. E., Peeters, M., Hahn, B. H., Ochman, H. (2016) Cospeciation of gut microbiota with hominids. Science 353(6297):380–382.

32. Sanders JG, et al. (2015) Baleen whales host a unique gut microbiome with similarities to both carnivores and herbivores. Nat Commun 6:8285.

33. Moeller AH, et al. (2017) Dispersal limitation promotes the diversification of the mammalian gut microbiota. Proc Natl Acad Sci U S A 114(52):13768– 13773.

34. Thomas T, et al. (2016) Diversity, structure and convergent evolution of the global sponge microbiome. Nat Commun 7:11870.

35. Avena CV, et al. (2016) Deconstructing the Bat Skin Microbiome: Influences of the Host and the Environment. Front Microbiol 7:1753.

36. Muletz Wolz CR, Yarwood SA, Campbell Grant EH, Fleischer RC, & Lips KR (2018) Effects of host species and environment on the skin microbiome of Plethodontid salamanders. J Anim Ecol 87(2):341–353.

37. Bryant JA, Lamanna, C., Morlon, H., Kerkhoff, A. J., Enquist, B. J., Green, J. L. (2008) Microbes and mountainsides: contrasting elevational patterns of bacterial and plan diversity. PNAS 105:11505–11511.

38. Singh D, et al. (2014) Strong elevational trends in soil bacterial community composition on Mt. Halla, South Korea. Soil Biology and Biochemistry 68:140– 149.

39. Carrillo-Araujo M, et al. (2015) Phyllostomid bat microbiome composition is associated to host phylogeny and feeding strategies. Front Microbiol 6:447.

40. Phillips CD, et al. (2012) Microbiome analysis among bats describes influences of host phylogeny, life history, physiology and geography. Mol Ecol 21(11):2617–2627.

41. Eckburg PB, Bik, E. M., Bernstein, C. N., Purdom, E., Dethlefsen, L., Sargent, M., Gill, S. R., Nelson, K. E., Relman, D. A. (2005) Diversity of human intestinal microbial flora. Science 308(5728):1635–1638.

42. Yildirim S, et al. (2010) Characterization of the fecal microbiome from non-human wild primates reveals species specific microbial communities. PLoS One 5(11):e13963.

43. Bik EM, et al. (2016) Marine mammals harbor unique microbiotas shaped by and yet distinct from the sea. Nat Commun 7:10516.

44. Gutman N, Hansen MJ, Bertelsen MF, & Bojesen AM (2016) Pasteurellaceae bacteria from the oral cavity of Tasmanian devils (Sarcophilus Harrisii) show high minimum inhibitory concentration values towards aminoglycosides and clindamycin. Lett Appl Microbiol 62(3):237–242.

45. Verhulst NO, et al. (2011) Composition of human skin microbiota affects attractiveness to malaria mosquitoes. PLoS One 6(12):e28991.

46. Busula AO, Takken W, JG DEB, Mukabana WR, & Verhulst NO (2017) Variation in host preferences of malaria mosquitoes is mediated by skin bacterial volatiles. Med Vet Entomol 31(3):320–326.

47. Verhulst NO, et al. (2009) Cultured skin microbiota attracts malaria mosquitoes. Malar J 8:302.

48. Robinson A, et al. (2018) Plasmodium-associated changes in human odor attract mosquitoes. Proc Natl Acad Sci U S A 115(18):E4209–E4218.

49. De Moraes CM, et al. (2014) Malaria-induced changes in host odors enhance mosquito attraction. Proc Natl Acad Sci U S A 111(30):11079–11084.

50. de Boer JG, et al. (2017) Odours of Plasmodium falciparum-infected participants influence mosquito-host interactions. Sci Rep 7(1):9283.

51. Song SJ, Amir, A., Metcalf, J. L., Amato, K. R., Xu, Z. Z., Humphrey, G., Knight, R. (2016) Preservation methods differ in fecal microbiome stability, affecting suitability for field studies. mSystems 1(3):1 – 12.

52. Caporaso JG, Lauber, C. L., Walters, W. A., Berg-Lyons, D., Lozupone, C. A., Turnbaugh, P. J., Fierer, N., Knight, R. (2011) Global patterns of 16S rRNA diversity at a depth of millions of sequences per sample. PNAS 108:4516– 4522.

53. Caporaso JG, et al. (2012) Ultra-high-throughput microbial community analysis on the Illumina HiSeq and MiSeq platforms. ISME J 6(8):1621–1624.

54. Kozich JJ, Westcott, S. L., Baxter, N. T., Highlander, S. K., Schloss, P. D. (2013) Development of a dual-index sequencing strategy and curation pipeline for analyzing amplicon sequence data on teh MiSeq Illumina sequencing platform. Applied and Environmental Microbiology 79(17):5122–5120.

55. Caporaso JG, Kuczynski, J., Stombaugh, J., Bittinger, K., Bushman, F. D., Costello, E. K., Fierer, N., Gonzalez Peña, A., Goodrich, E. K., Gordon, J. I., Huttley, G. A., Kelley, S. T., Knights, D., Koenig, J. E., Ley, R. E., Lozupone, C. A., McDonald, D., Muegge, B. D., Pirrung, M., Reeder, J., Sevinsky, J. R., Turnbaugh, P. J., Walters, W. A., Widmann, J., Yatsunenko, T., Zaneveld, J., Knight, R. (2010) QIIME allows analysis of high-throughput community sequencing data. Nature Methods 7(5):335–336.

56. Rognes T, Flouri T, Nichols B, Quince C, & Mahe F (2016) VSEARCH: a versatile open source tool for metagenomics. PeerJ 4:e2584.

57. McMurdie PJ, Holmes, S. (2014) Waste not, want not: Why rarefying microbiome data is inadmissable. PLoS Comput Biol 10(4):1–12.

58. McMurdie PJ & Holmes S (2013) phyloseq: an R package for reproducible interactive analysis and graphics of microbiome census data. PLoS One 8(4):e61217.

59. Oksanen J., et al. (2018) vegan: Community Ecology Package), R package version 2.5-2.

60. Wickham H (2016) ggplot2: Elegant Graphics for Data Analysis (Springer-Verlag, New York, NY).

61. Wickham H, François R, Henry L, & Müller K (2018) dplyr: A Grammar of Data Manipulation), R package version 0.7.6.

62. Breiman L (2001) Random Forests. Machine Learning 45:5–32.

63. Kurtz ZD, et al. (2015) Sparse and compositionally robust inference of microbial ecological networks. PLoS Comput Biol 11(5):e1004226.

64. Cardona C (2017) CAVNet: Creation Analysis and Visualization of Networks).

65. Csardi G, Nepusz, T. (2006) The Igraph Software Package for Complex Network Research. InterJournal, Complex Systems 1695.

66. Love MI, Huber W, & Anders S (2014) Moderated estimation of fold change and dispersion for RNA-seq data with DESeq2. Genome Biol 15(12):550.

